# Qualitatively different delay-dependent working memory distortions in people with schizophrenia and healthy control subjects

**DOI:** 10.1101/2023.04.04.535597

**Authors:** Sonia Bansal, Gi-Yeul Bae, Benjamin M Robinson, Jenna Dutterer, Britta Hahn, Steven J Luck, James M Gold

## Abstract

**Background:** Impairments in working memory(WM) have been well-documented in people with schizophrenia(PSZ). However, these *quantitative* WM impairments can often be explained by nonspecific factors, such as impaired goal maintenance. Here, we used a spatial orientation delayed-response task to explore a *qualitative* difference in WM dynamics between PSZ and healthy control subjects(HCS). Specifically, we took advantage of the discovery that WM representations may drift either toward or away from previous-trial targets(serial dependence). We tested the hypothesis that WM representations drift toward the previous-trial target in HCS but away from the previous-trial target in PSZ.

**Methods:** We assessed serial dependence in PSZ(N=31) and HCS(N=25), using orientation as the to-be-remembered feature and memory delays from 0 to 8s. Participants were asked to remember the orientation of a teardrop-shaped object and reproduce the orientation after a varying delay period.

**Results:** Consistent with prior studies, we found that current-trial memory representations were less precise in PSZ than in HCS. We also found that WM for the current-trial orientation drifted *toward* the previous-trial orientation in HCS(representational attraction) but drifted *away* from the previous-trial orientation in PSZ(representational repulsion).

**Conclusions:** These results demonstrate a qualitative difference in WM dynamics between PSZ and HCS that cannot easily be explained by nuisance factors such as reduced effort. Most computational neuroscience models also fail to explain these results, because they maintain information solely by means of sustained neural firing, which does not extend across trials. The results suggest a fundamental difference between PSZ and HCS in longer-term memory mechanisms that persist across trials, such as short-term potentiation and neuronal adaptation.

## INTRODUCTION

Since Park and Holzman(1), studies of working memory (WM) have shown that people with schizophrenia (PSZ) have significant deficits when asked to recall a spatial location after a delay period(2). In some (but not all) studies, deficits are present at short delays and increase as the delay interval increases, suggesting impairments in both encoding and maintenance processes(3–7).. Additionally, several functional magnetic resonance imaging studies report that PSZ exhibit abnormal activity in the brain areas associated with WM, such as prefrontal and parietal cortex (8–10).

A limitation of this research has been that *quantitative* impairments in behavioral performance can be explained by extraneous factors such as poor task comprehension, reduced motivation, and lapses of attention(e.g.(7,11–15)). In the present study, we focus on a *qualitative* difference between PSZ and healthy control subjects(HCS), in which opposite-direction WM distortions emerge across groups as the delay interval increases.

This research is based on a phenomenon from the cognitive science literature called *serial dependence*, in which memory for the target on trial N is unconsciously biased by the target from trial N-1. Depending on the conditions, the representation of the current-trial target is either attracted toward or repelled away from the previous-trial target(16–18). This phenomenon has been observed for a broad range of visual stimuli—including both simple features (e.g., orientation and spatial location) and more complex objects (e.g., faces)—and is presumably an automatic bias given that the target from trial N-1 is no longer task-relevant during trial N (19– 22). This carryover from previous trials implies a form of information storage that has not been previously considered in the clinical WM literature.

Both empirical data and computational models suggest that WM representations are maintained by sustained firing of neurons that represent the feature value of the target stimulus(23–25). This sustained firing terminates after the response is made to the current trial and cannot easily explain biases from the previous trial. These serial biases may instead be mediated by short-term synaptic plasticity mechanisms(26–28), which allow information to be maintained in synaptic weights in the absence of sustained neural activity. The short-term weight changes that occur on trial N-1 will bias the representation of the target on trial N, causing it to drift toward the N-1 target. As the current-trial delay increases, more drift will accumulate, causing greater attraction at longer delays.

Recently, Stein et al.(29) found that serial dependence over increasing retention periods is reduced in PSZ and people with anti-NMDAR encephalitis compared to HCS. The reported location of the current-trial target was *attracted* toward the previous-trial target in HCS, and this bias increased as the current-trial delay period increased from 0 to 4 seconds. This attraction was reduced in the encephalitis group, and it was actually reversed in PSZ, who showed suggestive evidence of delay-dependent *repulsion* of the current-trial representation away from the previous-trial location. Using a microcircuit model of the prefrontal cortex, Stein et al. found that altering the mechanisms that produce sustained firing during the current-trial delay period (e.g. the excitatory-inhibitory balance) could not account for the pattern of results. However, reduced NMDAR-dependent short-term plasticity(STP) could explain reduced delay-dependent attraction in both PSZ and people with anti-NMDAR encephalitis, and is consistent with prior studies suggesting reduced plasticity in PSZ(30–35). More broadly, these serial dependence effects may provide a behavioral means of assessing the integrity of short-term plasticity mechanisms.

The serial dependence findings of Stein et al.(29) are rare example of a striking qualitative cognitive performance difference between PSZ and HCS, with cross-trial attraction in HCS and cross-trial repulsion in PSZ. Because of the potential importance of qualitative differences, the present study sought to replicate and extend the findings of Stein et al. We had six goals. First, because Stein et al. tested only 17 PSZ, we wanted to replicate the qualitative difference using a larger sample. Second, we doubled the maximal WM delay interval to more clearly assess whether the group differences were amplified at longer delays. This delay amplification provides critical evidence that constrains the set of possible neural mechanisms. Third, we sought to examine the generalizability of their findings by using an orientation WM task rather than a spatial location WM task. Orientation WM is mediated largely by visual cortex rather than PFC (36–38), and the finding of a similar effect for orientation would suggest a cortex-wide mechanism rather than a PFC-specific mechanism. Fourth, we tested the hypothesis that greater drift in WM precision in PSZ relative to HCS reflects systematic effects of serial biases rather than the general instability of WM representations in PSZ postulated by previous models (39). Fifth, we asked whether the serial dependence in PSZ is correlated with within-trial repulsion effects that we previously observed when two orientations must be simultaneously maintained in WM on each trial (40). If this within-trial effect reflects the same mechanism as the across-trial serial dependence, then the two effects should be correlated, and this would further constrain the set of possible neural mechanisms. Finally, we assessed correlations with other measures of WM to determine whether the unusual pattern of serial dependence exhibited by PSZ is related to more conventional measures of WM dysfunction.

More broadly we hope to establish serial dependence as a robust means of obtaining qualitative rather than merely quantitative differences in behavior between PSZ and HCS.

## METHODS AND MATERIALS

### Participants

31 people with a diagnosis of schizophrenia or schizoaffective disorder (referred to as PSZ hereafter) were recruited from the Maryland Psychiatric Research Center (MPRC) and other local clinics. Diagnosis was established combining information from a Structured Clinical Interview for DSM-IV (SCID) with a review of medical records at a consensus diagnosis meeting. All PSZ were clinically stable outpatients who had been receiving the same antipsychotics, at the same dose, for at least 4 weeks prior to study participation. Healthy control subjects (HCS, N=25) were recruited via online advertisements and local bulletin boards. They were screened using the Structured Clinical Interview for DSM-IV-TR Axis I Disorders and the Structured Interview for DSM-III-R Personality Disorders–Revised. All HCS had no current Axis I disorder or Axis II schizophrenia spectrum disorder, neurological disorder, or cognitively impairing medical disorder, with no lifetime history of psychosis or history of psychotic disorders in first-degree relatives. After a complete study description was provided to all subjects, written informed consent was obtained. The protocol was approved by the Institutional Review Board of the University of Maryland, Baltimore. Demographic and clinical information is summarized in Table 1.

### Clinical and Neurocognitive Measures

In PSZ, symptom assessments included the Brief Psychiatric Rating Scale (BPRS;(41)) and the Scale for the Assessment of Negative Symptoms (SANS;(42)). All participants received the Measurement and Treatment Research to Improve Cognition in Schizophrenia (MATRICS) Consensus Cognitive Battery (43,44).

### Apparatus

Stimuli were generated in MATLAB (The MathWorks, Inc.) using Psych Toolbox (45,46) and were displayed on a LCD monitor with a gray background at a distance of 100 cm. Manual responses were collected using a computer mouse.

### Design and Procedure

In this *delayed estimation* WM task (see Figure 1A), participants had to remember the orientation of a teardrop-shaped object and reproduce the orientation after a varying delay period (0, 2, 4 or 8 s). A black fixation dot was continuously present in the center of the display except when occluded by the teardrop, and each trial began with a 1200-ms fixation interval. Each target was a teardrop shape (3° long, 1° maximum width) presented at the center of the display for 200 ms. The orientation of a given target was selected with equal likelihood from a set of 12 equally spaced values (separated by 30°, starting at 15° from upright to avoid horizontal and vertical orientations). Orientations from this set were tested in random order with equal probability. Although 12 discrete orientation values were used, this was not obvious to the participants. Participants were instructed to remember the orientation as precisely as possible over the delay period, during which only the fixation dot was visible.

**FIGURE 1.**
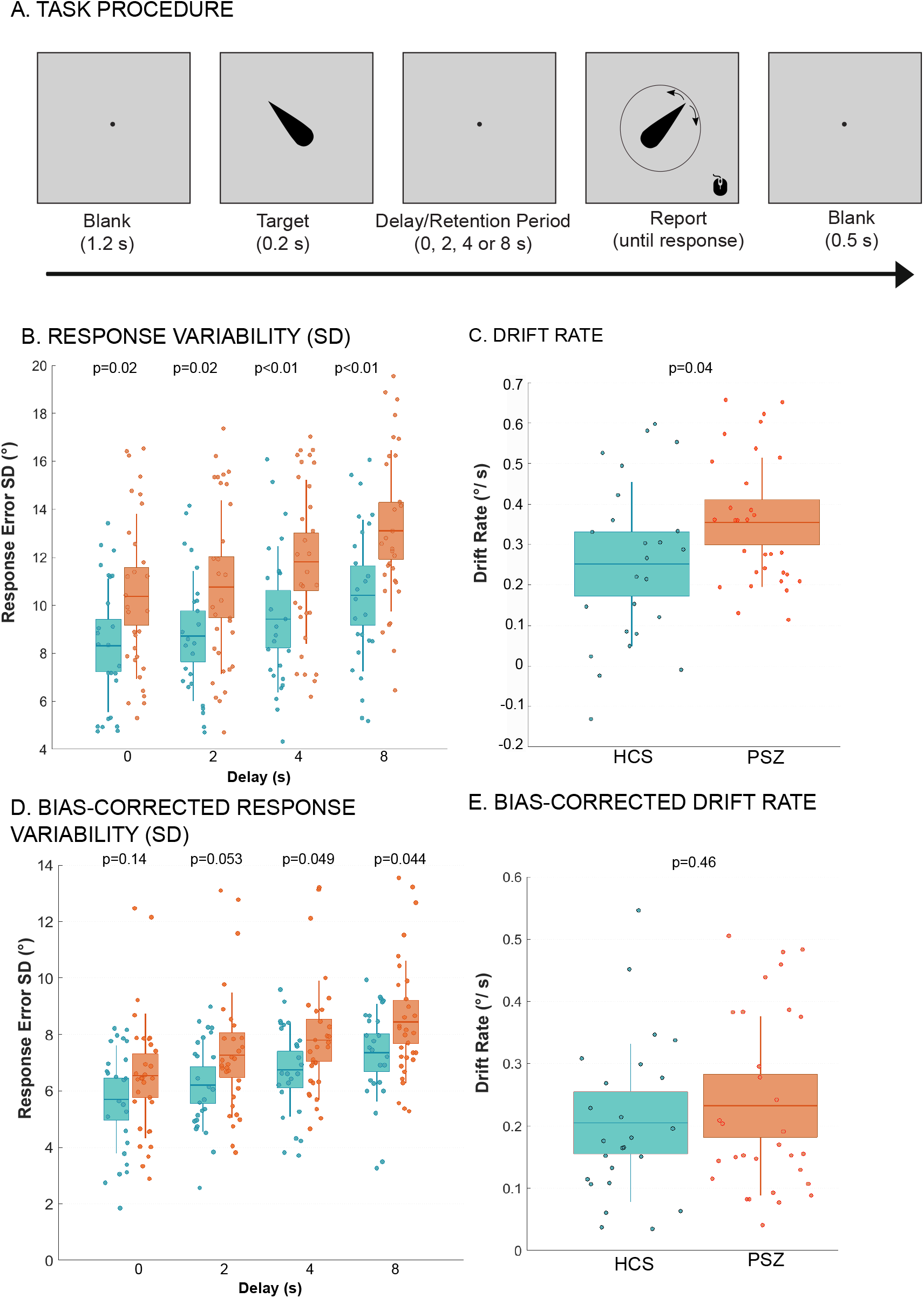
**A. Delayed estimation working memory task.** Participants were asked to remember the orientation of a teardrop-shaped object and reproduce the orientation after a varying delay period. The orientation of a given target was selected with equal likelihood from a set of 12 equally spaced values. Each trial began with a 1.2 s fixation dot, followed by a teardrop object presented at the center for 0.2 s. This was followed by a delay of 0, 2, 4, or 8 s, during which only the fixation dot was visible. At the end of the delay, participants reproduced the orientation of the target using a computer mouse. This report was followed by a 0.5 s intertrial interval. **B. Response Variability**. Average standard deviation of response errors. This variability (inverse precision) increased in magnitude over time. **C. Drift Rate**. This was calculated as the slope of the function relating standard deviation to delay period. **D**. Bias-corrected Response Variability. **E**. Bias-corrected Drift Rate.

At the end of the delay, participants reproduced the orientation of the target using a computer mouse. The mouse pointer started at the fixation point; once the pointer moved, a teardrop shape appeared at an orientation that matched the current position of the mouse. The participant adjusted the mouse position until the teardrop matched the remembered orientation of the target and then pressed the mouse button to finalize the response. We have previously shown that motor variability in this kind of mouse clicking task is near zero in both PSZ and HCS (44). The response was followed by a 500-ms intertrial interval. After 16 practice trials, each participant completed three blocks of 96 trials (total of 288 trials, 72 at each delay). The 12 orientations and 4 delay periods occurred in random order with equal probability.

### Analysis

The main question of the present study was whether the reported orientation on a given trial was influenced by the orientation on the previous trial (even though the orientations were presented in random order such that the previous-trial orientation did not predict the current-trial orientation). To answer this question, we examined the reported orientation on the current trial as a function of the previous-trial orientation, excluding the first trial in each block. With 12 orientations, there were 12 discrete differences between the current and previous-trial orientations. We collapsed the data across mirror-image orientation differences, producing seven different absolute orientation differences (0°, 30°, 60°, 90°, 120°, 150°, 180°). Data from trials with a 0° or 180° orientation difference were excluded from analysis because attraction and repulsion are not defined for these differences.

Our main dependent variable was the *response error*, defined as the difference between the reported orientation and the true orientation for the current trial. We coded the sign of this difference so that the response error reflected the bias relative to the previous-trial orientation: Positive values indicated that the current-trial report was biased towards the previous-trial orientation (attraction), whereas negative values indicated a bias away from the previous-trial orientation (repulsion). The response errors were averaged for each of the seven orientation differences (excluding orientation differences of 0° or 180°). We also excluded trials on which the current-trial response error was larger than 60° or the response initiation time exceeded 3 s, indicating a likely lapse of attention. This criterion removed 10.6±5.0% of trials in PSZ and 6.4±1.6% of trials in HCS (t=3.96, p=0.0002), suggesting PSZ were more prone to attentional lapses than HCS.

Precision for each subject and delay was estimated as the standard deviation (SD) of the absolute angular response error across trials to assess the trial-by-trial memory variability (i.e., the imprecision of the memory). We also quantified the slope of the function relating the SD to delay duration, which provides a measure of the WM drift rate.

The mean response error and precision measures for the individual participants were analyzed in a two-way ANOVA with factors of group (PSZ vs. HCS) and absolute orientation differences (five levels), with the Greenhouse-Geisser epsilon correction for nonsphericity.

### Within-Trial Repulsion

We also examined within-trial repulsion, using the paradigm described by Bansal et al. (37), but with one-fourth the amount of trials. Briefly, two teardrops were presented sequentially at the beginning of each trial (200 ms duration, separated by 750 ms), and participants were required to remember both teardrops simultaneously. After a 1000-ms delay interval, they reproduced the orientations of both teardrops (in a randomized order). Response errors were coded with respect to the other teardrop on the same trial.

## RESULTS

Background demographic and cognitive features are shown in Table 1. The groups were similar in age, gender, and ethnicity distributions and had similar parental education.

### WM Precision

We first assessed whether WM representations were less precise (i.e., more spatially variable) in PSZ than in HCS and whether this loss of precision was amplified over increasing delays. Response variability increased with delay duration in both groups (Figure 1B), leading to a significant main effect of delay duration (F_[2.52,136.26]_ = 63.81, p<0.001 η^2^_p_=0.54). PSZ had greater overall variability than HCS at all delay durations (significant main effect of group, F_[1,54]_ = 8.17, p=0.006, η^2^_p_=0.13). The increase in variability over delay durations was numerically greater for PSZ than for HCS, but this difference did not reach significance (delay duration × group interaction, F_[2.52,136.26]_ = 2.26, p=0.095,η^2^_p_=0.04). To provide a more sensitive metric of WM drift, we also conducted an analysis of the slope of the SD increase over time (Drift Rate, Figure 1C). The slope in PSZ (0.36° per second) was significantly larger than the slope in HCS (0.26° per second) (t = 2.12, p=0.039, Cohen’s d = 0.57). Post-hoc between-group comparisons for each delay duration are reported in Figure 1B. Note, we will later show that the difference in drift rate may be a secondary consequence of between-group differences in serial dependence.

### Effect of Delay Duration on Serial Dependence

We next assessed whether the reported orientation on the current trial was biased toward or away from the previous-trial, and whether this bias was influenced by the current-trial delay duration. Figure 2A displays the response bias as a function of the difference in orientation between the previous and current trial for the 0, 2, 4, and 8s delays. First consider the 8s delay. When the orientation difference between the previous trial and the current trial was smaller than 90°, the reported orientation was strongly biased towards the previous-trial orientation in HCS (attraction), whereas the reported orientation was strongly biased away from the previous-trial orientation in PSZ (repulsion). These attraction and repulsion effects were robust but somewhat smaller at the 4 s delay, declined further at the 2s delay, and disappeared at the 0s delay.

**FIGURE 2.**
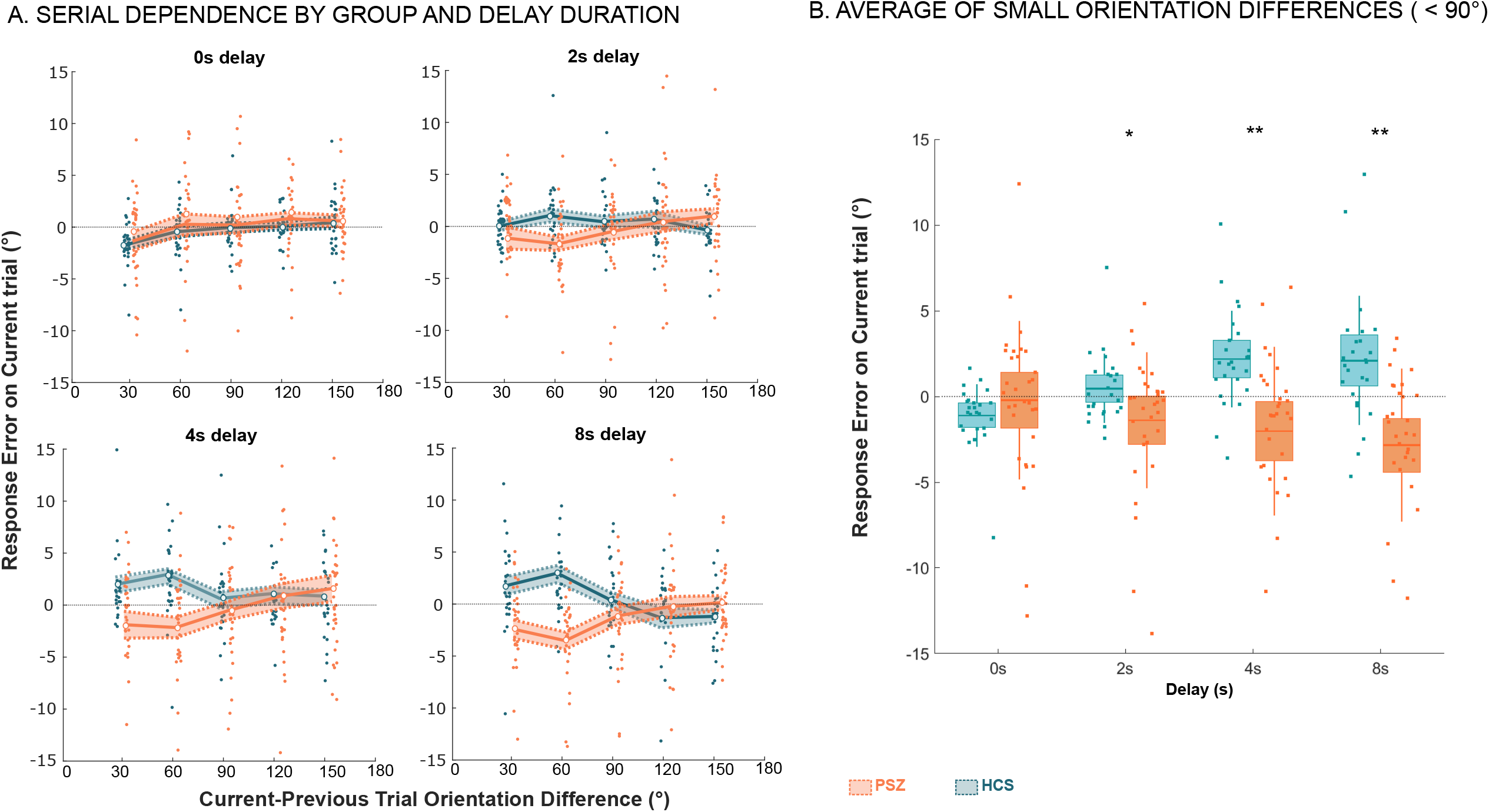
**A. Serial dependence by group and delay duration.** Serial dependence is calculated as the mean response error as a function of the difference in orientation between the previous and current trial. The single-subject means and the group means are plotted for each delay. Solid lines indicate the means, with shading being ± s.e.m. Dots indicate single-subject means. **B. Small Orientation Differences**. The mean bias index (mean response error averaged over trials with orientation differences of ≤90°) is plotted for each group (dots indicate single-subject means) at each delay, showing that the bias became progressively more positive over delays in HCS and progressively more negative over delays in PSZ.

These data were analyzed in a 3-way ANOVA with factors of group, orientation difference, and delay duration. The opposite direction of the biases in PSZ and HCS led to a significant Group X Orientation difference interaction (F_[4,216]_ = 7.23, p<0.001, η^2^_p_=0.12), but no main effect of Orientation difference (F_[4,216]_ =1.80, p=0.129, η^2^_p_=0.03). The delay dependence of this pattern led to a significant Group X Delay X Orientation difference interaction (F_[12,648]_ = 1.92, p=0.03, η^2^_p_=0.03).

Following Bae & Luck (22,47), we simplified the analysis by computing a single *bias index* for each participant, averaged over the trials with orientation differences of ≤90°. Figure 2B displays the mean bias index for each group at each delay, showing that the bias became progressively more positive over delays in HCS (attraction) and progressively more negative over delays in PSZ (repulsion). In a 2-way ANOVA with factors of Group and Delay, this pattern yielded a significant Group X Delay interaction (F_[3,162]_ = 2.79, p=0.043, η^2^_p_=0.05). As shown in Figure 2B, post-hoc between-group comparisons were significant for the 2, 4, and 8 s delays. One-sample *t* tests against zero showed that, at the 4 and 8 s delays, the attraction was significantly different from zero for HCS (4s delay: t=3.86, p<0.001, Cohen’s d=0.77, 8 s delay: t=2.78, p=0.01, Cohen’s d=0.56) and the repulsion was significantly different from zero for PSZ (4s delay: t= -2.28, p=0.03, Cohen’s d= -0.41, 8 s delay: t= -3.55, p<0.001, Cohen’s d= -0.64). Averaging across the 4 and 8 s delays, 80% of HCS showed attraction (positive values), whereas 84 % of PSZ showed repulsion (negative values).

We also examined whether the serial dependence varied according to the previous-trial delay period. In a 3-way ANOVA with factors of group, orientation difference, and previous trial-delay duration, no main effects or interactions involving the previous-trial delay were significant (η^2^_p_<0.03 in every case). This mirrors previous research in healthy young adults in which the current-trial delay but not the previous-trial delay impacted serial dependence (17).

### Factoring Out Serial Dependence from the Precision Measure

Previous research has proposed that WM representations are less stable in PSZ, leading to greater random drift as the delay interval increases (4–7). This is quantified by examining the trial-to-trial variability in responses at each delay (as in Figure 1B) and calculating the slope of the increase over delays (as in Figure 1C). However, rather than reflecting random drift, these effects could be a result of serial dependence (i.e., systematic drifts). It is therefore important to factor out the serial dependence effects when examining drifts in memory precision (see (29)).

To accomplish this, we recomputed the precision and drift measures shown in Figure 1 after correcting for serial dependence (see supplemental materials for details). Once the serial dependence bias was removed, we found that PSZ still had numerically poorer precision (larger SDs) than HCS at all delay durations, but this effect was no longer statistically significant in the omnibus ANOVA (main effect of group, F[1,54] = 3.82, p=0.06, η^2^p=0.07). Post-hoc tests revealed that the variability was significantly higher in PSZ for 4 and 8s delays (Figure 1B (p=0.049, p=0.044 respectively)). Thus, at least some of the apparent between-group difference in precision may be a result of systematic serial dependence rather than random errors.

When the slope of the corrected SD values was used to estimate the drift rate (Figure 1E), the corrected drift rate was similar for both groups (t=0.75, p=0.46, Cohen’s d=0.20). Thus, the greater drift rate reported in previous studies might be explained by systematic drifts toward or away from the previous-trial target—going in opposite directions for PSZ and HCS—rather than a general instability of WM in PSZ.

### Correlations with clinical and neurocognitive measures

We examined associations between the bias index and other measures of WM in both groups, as well as symptom ratings and medication dosage in PSZ. We collapsed the bias index across the 4s and 8s delays to obtain a unitary measure for correlations. The same bias index was computed for the within-trial repulsion task, in which two orientations were stored in WM on each trial. As shown in Figure 3, we found that the bias index from the current task was significantly correlated with the bias index from the within-trial task in PSZ (Spearman’s rho= 0.37, p=0.04), but not HCS (Spearman’s rho= 0.26, p=0.21). This suggests that the repulsive serial dependence observed at long delay periods in PSZ may reflect the same mechanism that causes simultaneous WM representations to repel each other.

**FIGURE 3.**
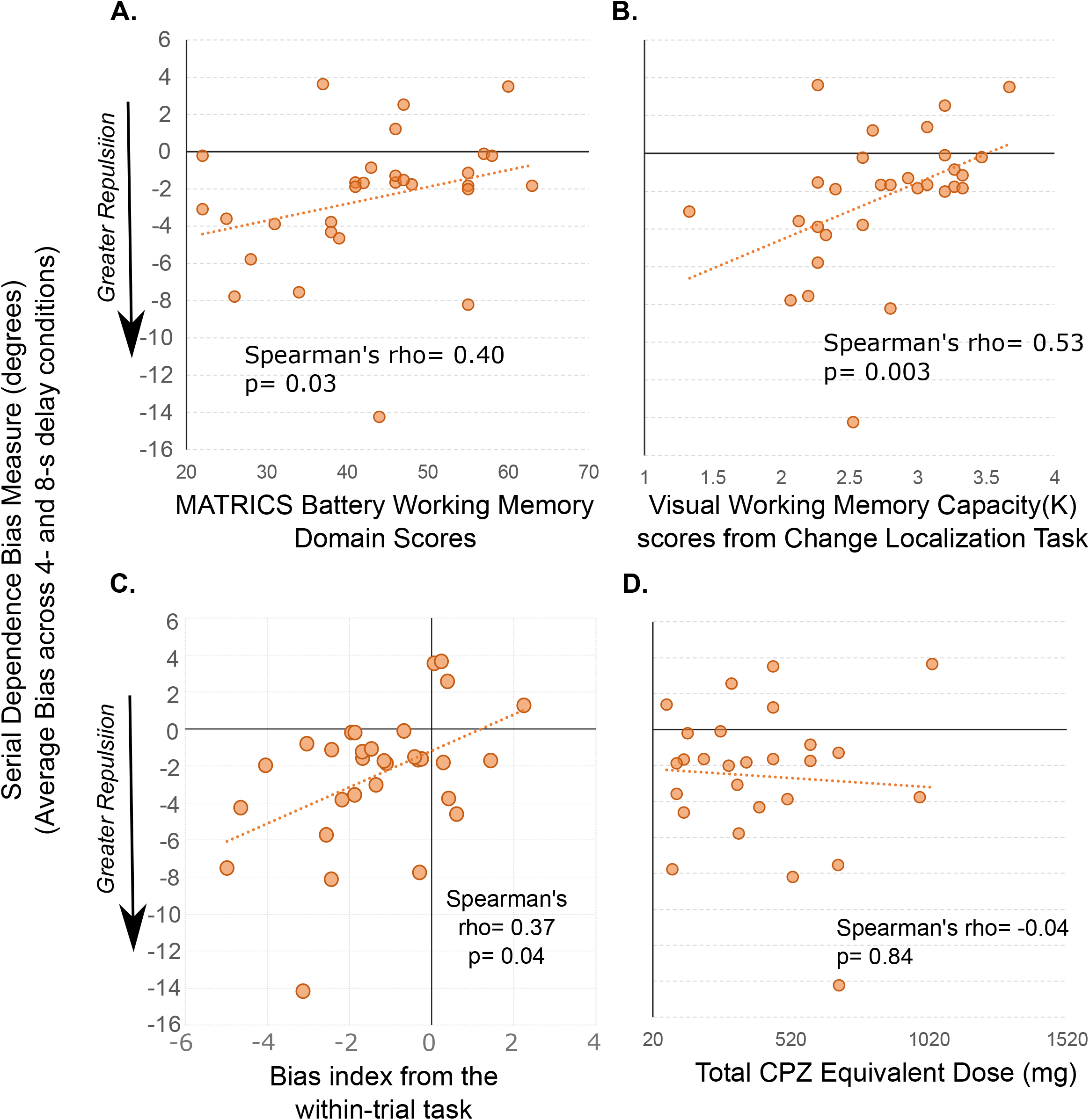
Correlations with neurocognitive and clinical measures in PSZ. **A**. In PSZ, serial dependence bias was associated with the working memory cognitive domain from the MATRICS battery. **B**. In PSZ, serial dependence bias was also associated with visual working memory capacity (K) from a change localization task. Greater repulsion was associated with lower scores on WM measures in PSZ. **C**. In PSZ, serial dependence bias in this task was correlated with the bias index from the within-trial task. **D**. Correlation between bias and medication dose (not significant).

Additionally, in PSZ, but not in HCS, we observed significant associations between serial dependence bias and the working memory cognitive domain from the MATRICS battery (PSZ, Spearman’s rho=0.53, p=0.003; HCS, Spearman’s rho=-0.17, p=45), and visual working memory capacity (K) from a change localization task (see Supplement for a description; PSZ, Spearman’s rho=0.40, p=0.03; HCS, Spearman’s rho=0.03, p=0.91). Thus, greater repulsion was associated with lower scores on WM measures in PSZ. We did not observe any significant correlations between bias and overall symptom (BPRS total, Spearman’s rho=-0.31 p=0.10), and there were no significant correlations with medication dose (Spearman’s rho=-0.04 p=0.84). Total MATRICS scores were not correlated with overall bias in either group (p>0.24).

## DISCUSSION

Whereas almost all previous studies of WM in PSZ found a *quantitative* impairment relative to HCS, Stein et al. (29) found a *qualitative* difference in WM biases produced by the previous-trial target, with an attractive bias in HCS and a suggestive repulsive bias in PSZ. This sort of opposite-direction, qualitative difference between PSZ and HCS is rare in schizophrenia research, and it is potentially important because it cannot easily be explained by nonspecific factors such as lapses of attention. However, this opposite-direction effect was a surprise to Stein and colleagues because it was not predicted by any known computational model of schizophrenia. Improbable effects are more likely to be spurious than expected effects, especially with modest sample sizes (48), so we sought to replicate this effect in a larger sample and determine whether it generalized to orientation memory. We found a robust, delay-dependent attractive bias in HCS and a robust, delay-dependent repulsive bias in PSZ, confirming that this qualitative difference between groups is replicable and generalizable. This effect may be an important clue about fundamental differences in cortical function in PSZ. We further showed that the degree of repulsion in PSZ is correlated with the degree of impairment in two different WM capacity measures.

Before discussing the mechanisms that may be implicated in these opposing biases, we would like to highlight how our WM precision results fit with the current WM literature. We expected that PSZ would show less precise WM than HCs, and that this loss of precision would be amplified with increasing the delay interval because the WM representations are assumed to be less stable in PSZ (2,4–7). This is precisely what we found: we observed that WM representations were less precise for longer delays in both groups, with a greater drift rate in PSZ than in HCS. However, when we factored out serial dependence, the drift rate was similar for both groups. These bias-corrected results suggested that previously published observations of greater WM drift rates in PSZ (including our own (4)) may be partially—or perhaps entirely— explained by these previously unknown differences in serial dependence. That is, PSZ may not exhibit greater *random* drifts in WM, as would be expected from a general instability, but may exhibit greater *systematic* drifts.

It is challenging to account for these findings mechanistically given how little work has been done on the neural basis of attractive and repulsive serial dependence effects. As noted earlier, Stein et al. (29) used computational modeling to demonstrate that alterations in STP, but not alterations in excitatory/inhibitory balance, can account for the reduced serial dependence seen in their NMDA encephalitis patients. However, reduced STP by itself cannot explain the delay-dependent repulsion observed in PSZ. Thus, an additional process must be responsible for this effect.

One plausible explanation of repulsive serial dependence is neuronal adaptation (49). For example, viewing a left-tilted orientation for a period of time will cause a subsequent vertical orientation to be perceived as slightly right-tilted (i.e., repelled away from the adapting stimulus). These aftereffects arise, in part, due to attenuation in the responses of neurons that code the features of the prior stimulus (feature-specific adaptation), which then biases the population response to subsequent stimuli away from the adapted features (50–52). There is growing evidence from the basic cognitive literature that serial dependence in WM reflects a mixture of repulsion arising from this sort of neuronal adaptation with attraction arising from STP or some other mechanism (53,54). That is, the presence of attraction versus repulsion in the behavioral responses depends on whether the adaptation-related repulsion is stronger or weaker than the STP-related attraction.

Thus, the repulsion observed in PSZ could reflect a massive reduction in STP, unmasking the repulsive effects of neuronal adaptation. Alternatively, STP could be equivalent in PSZ and HCS, but the adaptation could be greater in PSZ than in HCS, overwhelming the attractive bias. There is some evidence of greater adaptation effects in PSZ than in HCS for orientation and direction of motion (reviewed in (55)). Relatedly, we found that the repulsion effect exhibited by PSZ at long delays was correlated with amount of repulsion seen in a task where two orientations had to be maintained on each trial. We had previously found that this within-trial repulsion was greater in PSZ than in HCS and speculated this might reflect greater lateral inhibition between the WM representations in PSZ. Such a mechanism cannot explain the serial dependence observed across trials, but greater neuronal adaptation effects in PSZ could produce increased repulsive biases both between and within trials.

It will require programmatic experiments to distinguish between a reduction of STP or an increase in adaptation in PSZ. Given that these different alternatives reflect distinct neural mechanisms, such experiments would be very worthwhile. If the neural mechanisms can be pinpointed, then serial dependence may offer a powerful behavioral method to assess a marked alteration of cortical function in PSZ.

## ACKNOWLEDGEMENTS AND DISCLOSURES

This work was supported by the National Institute of Mental Health (Grant No.R01MH065034 [to JMG and SJL]). JMG receives personal fees from Acadia Pharmaceuticals Inc outside the submitted work and royalty payments from the Brief Assessment of Cognition in Schizophrenia. All other authors report no biomedical financial interests or potential conflicts of interest.

## TABLE CAPTION

**TABLE 1. Participant Characteristics, Cognitive and Clinical Measures**

