## Supplemental Material for "Qualitatively different delay-dependent working memory distortions in people with schizophrenia and healthy control subjects"


Contents Pg.


1. Correction for Serial Dependence Bias……………………………………………………2
2. Change Localization Task …………………………………………………………………3

### S1. **FACTORING OUT SERIAL DEPENDENCE BIAS**

Previous research has proposed that WM representations are less stable in PSZ, leading to greater random drift as the delay interval increases[1-4]. However, rather than reflecting true imprecision over time, these effects could potentially be a result of systematic biases. It is therefore important to factor out the serial dependence bias related to the previous trial stimulus when examining memory precision.

In order to factor out biased responses, and examine precision (inverse precision: circular standard deviation(SD)) related to the current trial's orientation stimulus, we followed the procedure below:

1. We categorized responses centered around the true changed direction. This was done by categorizing responses in 15° bins around the current trial orientation, T.
2. For these responses, we calculated response errors as the angular difference between the reported orientation and the true orientation for the current trial
3. Response errors were included in the calculation of response variability(SD) if
   - (i) they were centered in 15° bins around the true current trial orientation, T
   - (ii) they fell **outside** the demarcation of ±2 standard deviations around the previous trial (PT) orientation (i.e. outside the shaded pink region). Thus, in this example, responses in the green region were included, but not if they overlapped with the shaded pink regions.
   - (iii) Once this subset of responses centered around the true orientation of the current trial were identified, the mean standard deviation per delay was calculated.

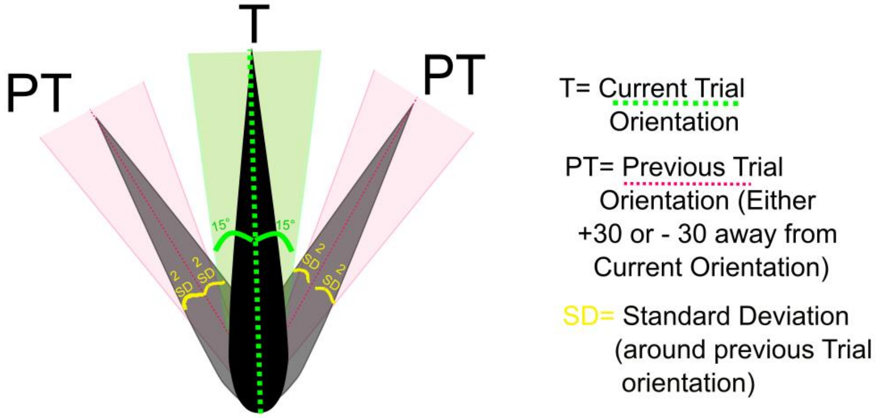


1. Gold JM, Bansal S, Anticevic A, Cho YT, Repovš G, Murray JD, et al. (2020): Refining the Empirical Constraints on Computational Models of Spatial Working Memory in Schizophrenia. Biol Psychiatry Cogn Neurosci Neuroimaging 5: 913–922.
2. Badcock JC, Badcock DR, Read C, Jablensky A (2008): Examining encoding imprecision in spatial working memory in schizophrenia. Schizophr Res 100: 144–152.
3. Mazhari S, Badcock JC, Waters FA, Dragović M, Badcock DR, Jablensky A (2010): Impaired spatial working memory maintenance in schizophrenia involves both spatial coordinates and spatial reference frames. Psychiatry Res 179: 253–8.
4. Starc M, Murray JD, Santamauro N, Savic A, Diehl C, Cho YT, et al. (2017): Schizophrenia is associated with a pattern of spatial working memory deficits consistent with cortical disinhibition. Schizophr Res 181: 107–116.

### S2. CHANGE LOCALIZATION TASK AND ANALYSIS

**Task[5]:** Participants completed 60 trials of a change localization task, an experimental paradigm that measures visual working memory (WM) capacity. The task was programmed in E-Prime, and stimuli were presented on an LCD monitor at a nominal viewing distance of 70 cm. As illustrated, a sample array of four colored squares, each measuring 0.7 × 0.7° of visual angle, for 100 ms was presented to participants. After a 900-ms delay during which only the fixation cross was shown, a test array was presented: This array was identical to the sample array except that one color had changed to a value that had not been present in the sample array. The colors were selected at random without replacement from a set of six colors with the following red, green, blue values: red (255, 0,0), green (0, 255, 0), yellow (255, 255, 0), magenta (255, 0, 255), cyan (0, 255, 255), and bright blue (80, 60, 255). From trial to trial, the colored squares were randomly placed around an imaginary circle with a radius of 3°, with the constraints that one item appeared in each of the 4 screen quadrants, and each item was separated by a minimum of 30° on the circle from the next item. Participants used a mouse to click on the location of the item in the test array that they believed had changed color from the sample array. Participants were encouraged to respond as accurately as possible, with no speed pressure, and a response was required on every trial. The test array was visible until the response was made, and trials were separated by a 2000-ms inter-trial interval. A total of 60 trials was tested in each subject, which typically required approximately 8–10 minutes of testing time.

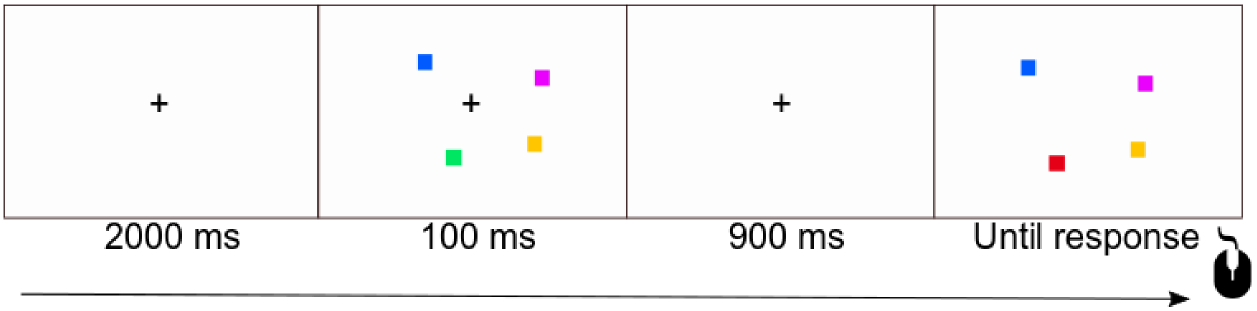


**Analysis:** WM storage capacity was quantified using a variant of the Pashler/Cowan K equation [6,7], where K signified how many items worth of information were stored in WM. Since each trial contained a change, there was no potential for false alarms. As a result, K was calculated by multiplying each participant's proportion correct by 4 (the number of items in the memory array).


5. Johnson, M. K., McMahon, R. P., Robinson, B. M., Harvey, A. N., Hahn, B., Leonard, C. J., ... & Gold, J. M. (2013). The relationship between working memory capacity and broad measures of cognitive ability in healthy adults and people with schizophrenia. *Neuropsychology*, *27*(2), 220.

6. Pashler H. Familiarity and visual change detection. *Perception and Psychophysics.* 1988; 44:369–378

7. Cowan N. The magical number 4 in short-term memory: A reconsideration of mental storage capacity. *Behavioral and Brain Sciences.* 2001; 24:87–185
